# Illuminating the dark side of the human transcriptome with TAMA Iso-Seq analysis

**DOI:** 10.1101/780015

**Authors:** Richard I. Kuo, Yuanyuan Cheng, Jacqueline Smith, Alan L. Archibald, David W. Burt

## Abstract

The human transcriptome is one of the most well-annotated of the eukaryotic species. However, limitations in technology biased discovery toward protein coding spliced genes. Accurate high throughput long read RNA sequencing now has the potential to investigate genes that were previously undetectable. Using our Transcriptome Annotation by Modular Algorithms (TAMA) tool kit to analyze the Pacific Bioscience Universal Human Reference RNA Sequel II Iso-Seq dataset, we discovered thousands of potential novel genes and identified challenges in both RNA preparation and long read data processing that have major implications for transcriptome annotation.

## Introduction

The transcriptome remains a vastly under explored space despite its significance as a foundation for biology. For eukaryotic species, alternative transcription start sites and RNA processing can result in a combinatorial array of transcript sequences which poses many challenges for understanding which RNA are functional and how they should be annotated^1^. Current genome annotations have mostly been based on data from low throughput cDNA sequencing and high throughput short read RNA sequencing^2^. However, cDNA sequencing cannot provide a reasonable coverage of the transcriptome and short read sequencing leads to issues with transcript model assemblies^3^.

The recent development of high throughput long read RNA sequencing promises a new age of transcriptome exploration^4^. Full length transcript reads provides high confidence in predicting alternative transcripts and distinguishing real transcripts from sequencing noise^5^.

While there have been many studies using long read RNA sequencing for transcriptome discovery^5,6,7,8,9^, the tools used for processing long read data suffer from limitations that severely reduce the sensitivity and specificity of transcriptome exploration. These strategies either rely on orthogonal information which biases gene discovery and are only available for a small number of species^10^ (Talon^11^, TAPIS^6^, SQANTI^12^) or on algorithms with serious theoretical limitations^13,14^.

The non-orthogonal strategies are based on inter-read error correction for removing errors from long read sequence data. These inter-read methods are split into two main strategies: long read error correction and short read error correction. Long read error correction involves the clustering of long reads from the same sequencing run or across multiple runs for error correction^15^. Pacific Biosciences’ Cluster/Polish method (previously known as Iterative Clustering for Error correction) is the most popular tool for doing long read error correction. In short read error correction, long reads are error corrected by aligning short read RNAseq data and replacing the long read sequences with the higher accuracy short read sequences^13,14,16^. Both long read and short read error correction methods have the drawback of the possible introduction of chimeric sequences due to the merging of sequence information across reads. This type of error occurs when the alignment of reads is compromised by regions of high error density thus allowing for reads from different transcripts to be grouped (Fig. 1). These methods are performed prior to mapping the reads to the genome.

**Figure 1.**
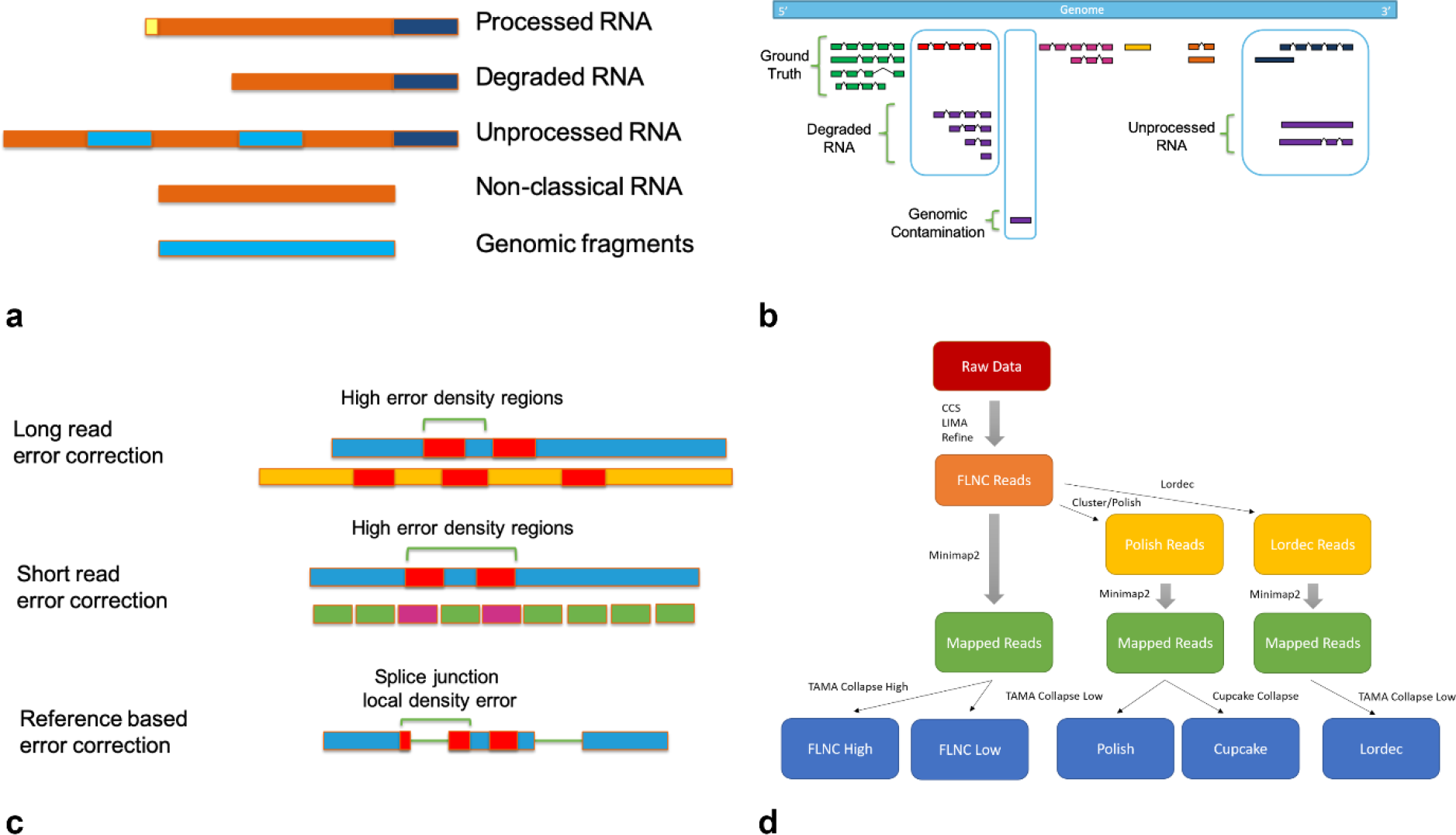
Long read RNA sequencing diagrams. (a) Types of sequences found in RNA samples. (b) Representation of RNA sample sequences relative to the genome. (c) Illustration of problems arising from different error correction methods for long reads. (d) Diagram of pipelines analyzed.

These inter-read correction strategies also do not fully account for reads originating from transcriptional noise. Within any RNA sample collected from a eukaryotic species, there is a mixture of mature functional RNA as well as pre-processed RNA, degraded RNA, and possible genomic contamination^12^ (Fig. 1). Differentiating between these sources of read information is crucial for producing high quality transcriptome annotations and preventing the false identification of novel genes.

To understand the extent of the limitations of currently available long read RNA analysis software, we analyzed the Universal Human Reference RNA (UHRR) Sequel II Iso-Seq data released by Pacific Biosciences (PacBio) using our Transcriptome Annotation by Modular Algorithms (TAMA) tool kit. TAMA is designed to take advantage of long read RNA data and high quality reference genome assemblies to produce the most accurate and informative transcript models theoretically possible given these inputs. This makes TAMA useful for situations where additional types of data are not available^17,18^. In addition, by not relying on orthogonal information, TAMA also provides a more agnostic approach to transcriptome annotation which can reveal problems with prior assumptions from previous annotation efforts.

## Results

### Pipeline sensitivity comparison

We processed the UHRR Iso-Seq data using 5 different pipelines to compare the effect of each method on gene discovery and model prediction accuracy. These pipelines include two pipelines without inter-read error correction (FLNC Low and FLNC High), two pipelines using long inter-read error correction (Polish and Cupcake), and one pipeline using short inter-read error correction (Lordec).

We first used a low stringency pipeline (FLNC Low) to estimate the upper limit of the number of possible gene and transcripts within the UHHR Iso-Seq data. This involved no pre-map error correction (mapping FLNC reads directly) and a TAMA Collapse run with low stringency parameters.

This FLNC Low pipeline resulted in 168,328 gene models with 752,996 transcript models (31,115 multi-exonic genes and 514,364 multi-exonic transcript models). The number of predicted genes and transcripts far exceeds the numbers found in the Ensembl human genome annotation v96^19^. These elevated numbers suggest the presence of a large amount of either sample noise or wet lab processing noise.

The large number of gene models were mostly made up of single exon genes (Fig. 2). Single exon genes can be produced from genomic contamination where adapters were able to prime either DNA fragments or RNA produced from run-on transcription^20^. The 3’ primer for current cDNA library preparation methods contains an oligo-dT region for capturing the poly-A tails of mature RNA. Since these primers only require a repeat of A’s to bind, they can bind to a genomic stretch of A’s thus introducing these sequences into the cDNA library^21^. To identify and filter out reads that may have originated from these events, we developed the tama_remove_polya_models_levels.py tool. This tool can be used to remove all transcript models with a genomic poly-A stretch on the 3’ end of the mapped read. Using tama_remove_polya_models_levels.py to remove all single exon models with 3’ genomic poly-A stretches resulted in 35,577 genes and 614,279 (24,133 multi-exonic genes and 507,382 multi-exonic transcripts) which equates to 21.1% of the genes and 81.6% of transcripts predicted in the FLNC Low pipeline.

**Figure 2.**
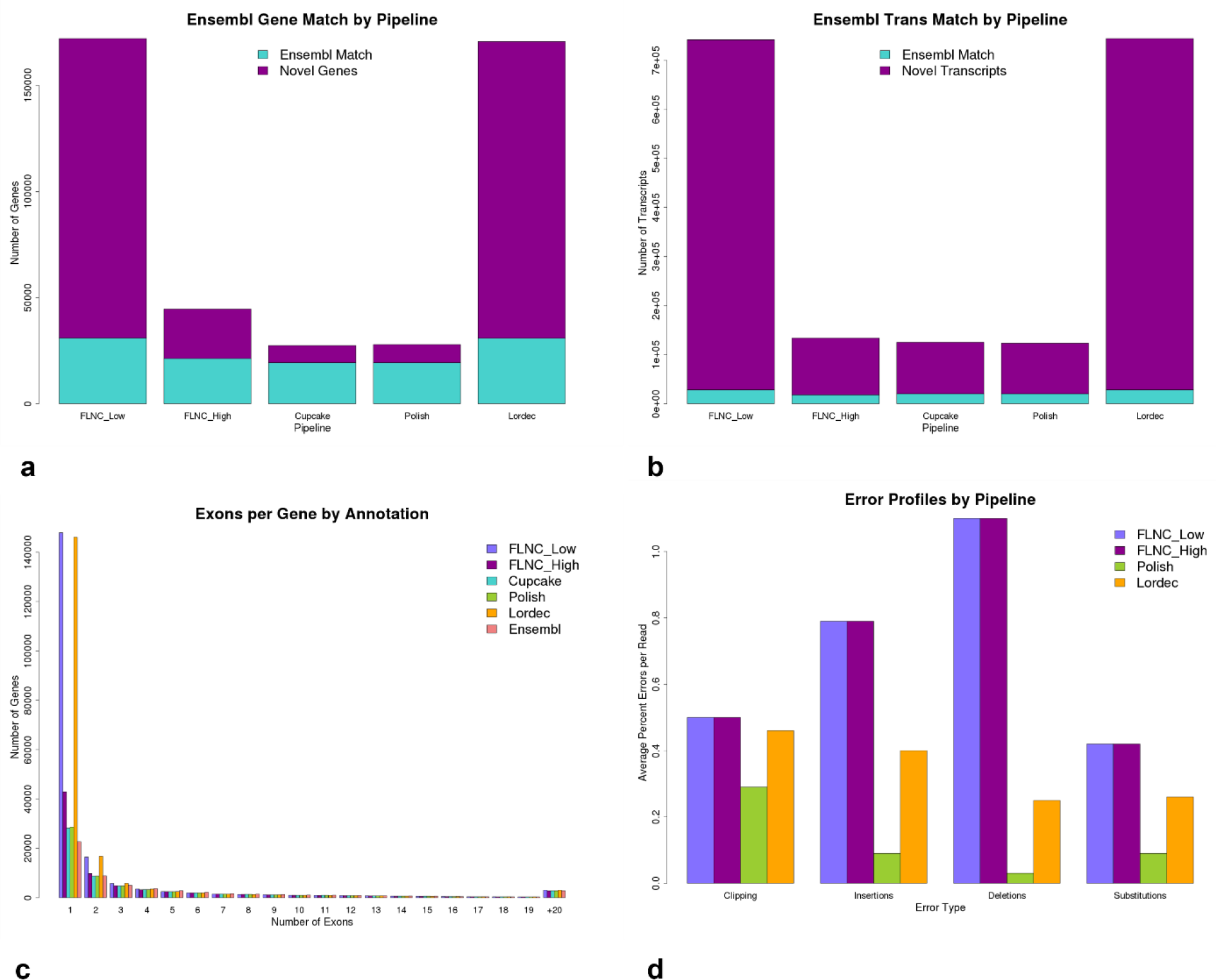
Comparing the results of the different Iso-Seq pipelines. (a) Number of novel and Ensembl matching genes by pipeline. (b) Number of novel and Ensembl matching transcripts by pipeline. (c) Maximum number of exons per gene by pipeline. (d). Type and amount of error per pipeline.

While the gene numbers are more reasonable after this filtration step, the transcript numbers still appear to be artificially high. This also suggests that the filtered genes (132,751) were likely the product of genomic contamination, noisy transcription in intergenic regions which span genomic poly-A regions, and/or transcribed processed pseudogenes. These filtered genes should be annotated if they are from transcribed processed pseudogenes since they would be technically part of the transcriptome and may have functionality^22^. However, if they are from genomic contamination and noisy transcription, this would suggest that the UHRR has a significant amount of contamination from either of those 2 sources. Since the UHRR is commonly used as a baseline for RNA experiments, this contamination would need to be handled bioinformatically to avoid erroneous interpretations.

To address the suspiciously large number of transcript models from our FLNC Low run, we ran a high stringency pipeline (FLNC High) with TAMA Collapse at a setting to remove transcripts with more than 1 error within a 20 bp range of a splice junction. We allowed for 1 bp of error due to possible true genomic variation between the UHHR samples and the reference genome assembly. We then then only kept transcript models with read support from both SMRT cells using tama_remove_single_read_models_levels.py. We required read support from both SMRT cells to avoid using reads that originated from PCR artefacts since PCR artefacts could be sequenced by multiple reads if their relative abundance is high enough. This assumes that the libraries for each SMRT cell were prepared separately. This high stringency pipeline resulted in 38,743 genes with 135,218 transcript models (15,777 multi-exonic genes and 87,112 multi-exonic transcripts) which equates to 23.0% of the genes and 18.0% of the transcripts predicted in the FLNC Low pipeline. We did not remove reads with genomic poly-A in this pipeline since real transcript models can have downstream genomic poly-A and the 2 SMRT cell support filtration is meant to remove PCR artefacts such as those caused by internal priming.

We compared both the FLNC Low and FLNC High annotations with the Ensembl v96 human annotation using TAMA merge to see how many models matched the public annotation (Fig. 2). We used two definitions for matching: gene level matches and transcript level matches. Gene level matches are defined as the number of genes with overlap between gene models (between annotations) on the same strand. Transcript level matches are defined as the number of transcripts that have the same exonic structures (between annotations) which is described in more detail in the methods section. The FLNC Low annotation had 30,947 gene level and 28,234 transcript level matches with Ensembl. However, only 13,874 genes had at least one transcript level match with the Ensembl annotation. The FLNC High annotation had 21,284 gene level and 17,932 transcript level matches with Ensembl with 10,649 genes that had at least one transcript level match.

Thus while the FLNC High pipeline produces a more reasonable number of genes and transcripts it still identifies 23,302 novel gene models. The FLNC Low pipeline identifies 9,663 more known genes as compared to the FLNC High pipeline. This suggests that choosing more stringent thresholds for filtering leads to a significant loss in real signal from the data.

We also compared the FLNC Low and FLNC High to other commonly used pipelines for Iso-Seq data. These include running the cluster/polish^15^ step for long read error correction (Polish pipeline), collapsing with Cupcake^15^ (Cupcake pipeline), and short read error correction with Lordec^13^ (Lordec pipeline).

For the Polish pipeline we ran the Isoseq3 cluster/polish tool on the FLNC reads and then processed the mapped reads with TAMA collapse using the same settings as the FLNC Low pipeline. This resulted in 25,731 genes and 126,288 transcripts (15,418 multi-exonic genes and 107,637 multi-exonic transcripts) (Fig. 2). The overall lower numbers of genes and transcripts is due to a filtration that occurs during the cluster/polish step which removes all reads that do not cluster with at least one other read. These non-clustered reads are called singletons.

For the Cupcake pipeline, we tested the effect of the collapsing algorithm by using the Cupcake collapse tool instead of TAMA Collapse as was used in the Polish pipeline. This resulted in 25,239 genes and 128,389 transcripts (15,395 multi-exonic genes and 110,200 multi-exonic transcripts) (Fig. 2). Thus the overall numbers were very similar between Cupcake collapse and TAMA collapse when working with clustered reads.

We ran the Lordec pipeline, to compare short read error correction. In the Lordec pipeline, we corrected the FLNC reads using Lordec with short read RNA data from the UHRR but from another study^23^. We then ran TAMA Collapse using the same settings as in the FLNC Low pipeline. This resulted in 166,766 genes and 753,756 transcripts (31,465 multi-exonic genes and 517,244 multi-exonic transcripts) (Fig. 2). These numbers are very similar to the FLNC Low pipeline results. Lordec does not discard or split reads as other hybrid correction tools do. Thus all reads are retained for mapping. This explains the similarity in numbers with the FLNC Low results.

When comparing the number of exons per gene across all 5 pipelines, the vast majority of the novel genes from the low stringency pipeline and the Lordec pipeline were single exonic (Fig. 2). Single exonic gene models are potentially suspicious as they could be the result of genomic contamination.

### Error comparison between pipelines

The number of genes and transcripts predicted from each pipeline provides some information about their respective sensitivity. However, other metrics need to be used to assess the accuracy of each pipeline. To understand how each pipeline dealt with errors, we looked at the error profiles for each mapped read for the FLNC Low, FLNC High, Polish, and Lordec pipelines. The Cupcake pipeline was omitted in this analysis because Cupcake does not provide a report on the errors in the mapped reads.

Using the output from TAMA Collapse we looked at length of coverage, identity, clipping, insertions, deletions, and substitution errors. These values represent the comparison of the mapped reads to the genome assembly and thus only serve as an estimate of the true rates of error.

Both FLNC Low and FLNC High pipelines had average coverage and identity values of 92.38% and 90.17% respectively. Note that the mapped FLNC reads are the same for these two pipelines thus all the error profiles of the mappings will be identical. The Polish pipeline produced an average coverage and identity of 97.84% and 97.53% respectively. While the Lordec pipeline had average coverage and identity of 92.44% and 91.48% respectively. Thus the Polish pipeline seems to out-perform the other pipelines in this metric. The Lordec pipeline values were unexpectedly similar to the FLNC pipelines suggesting that Lordec correction did not provide a large gain in error correction.

We then looked at overall error rates between the pipelines. To calculate average error rates, we counted the number of base pairs that were not matching between the mapped read and the genome sequence and divided this number by the length of the mapped read. This includes soft clipping, insertion, deletion, and substitution errors but does not include hard clipping. The FLNC Low and FLNC High pipelines have an average error rate of 2.83% per mapped read with an average mapped length of 1,959 bp. The Polish pipeline has an average error rate of 0.52% and average mapped read length of 2,218. The Lordec pipeline has an average error rate of 1.38% with a mapped read length average of 1,968 bp.

The longer average mapped read length of the Polish pipeline is most likely due to the cluster/polish algorithm merges read sequences with up to 100 bp length differences on the 5’ end. This behavior essentially absorbs the shorter reads into the longer reads effectively removing their length representation.

The Lordec pipeline had a similar amount of clipping errors as compared to the FLNC pipelines but lower rates of insertion, deletion, and substitution errors (Fig. 2). This indicates that the Lordec correction seem to be increasing the quality of the reads overall but has some issues correcting the ends of reads.

### Wobble comparison between pipelines

While the error rates of mapped reads are often used to assess the improvement of long read data from different pipelines^14^, this metric is actually not quite as useful for understanding the overall improvement in the transcriptome annotation. In genome based transcriptome annotations, the most important features to identify with respect to transcript models are the exact starts and ends of each exon as well as real exon chaining. Thus if the errors in the reads do not affect the overall transcript model, then there is no appreciable difference between a read with 10% error rate versus one with 0.1% error rate.

We measured this aspect of transcriptome annotation improvement by comparing the wobble at splice junctions with respect to transcript models annotated in the Ensembl human annotation. Wobble refers to small differences in mapped exon starts and ends (Fig. 3). So while two transcripts can have nearly identical structures, there can be small differences between their exon starts and ends which can be challenging to resolve. For the definition of how we defined “nearly identical” structure, see the methods section. Wobble typically occurs due to the higher error rate of long read sequences leading to small shifts in mapping the ends of each exon^24^. The amount of wobble between the transcript models of each pipeline compared to the reference annotation, provides a metric for the actual differences in the final transcriptome annotations produced by each pipeline. We ignored wobble at the transcript start and end sites due to the high variance of these features in natural RNA^25,26^. We also only assessed Ensembl transcript models that had coverage from all assessed pipelines.

**Figure 3.**
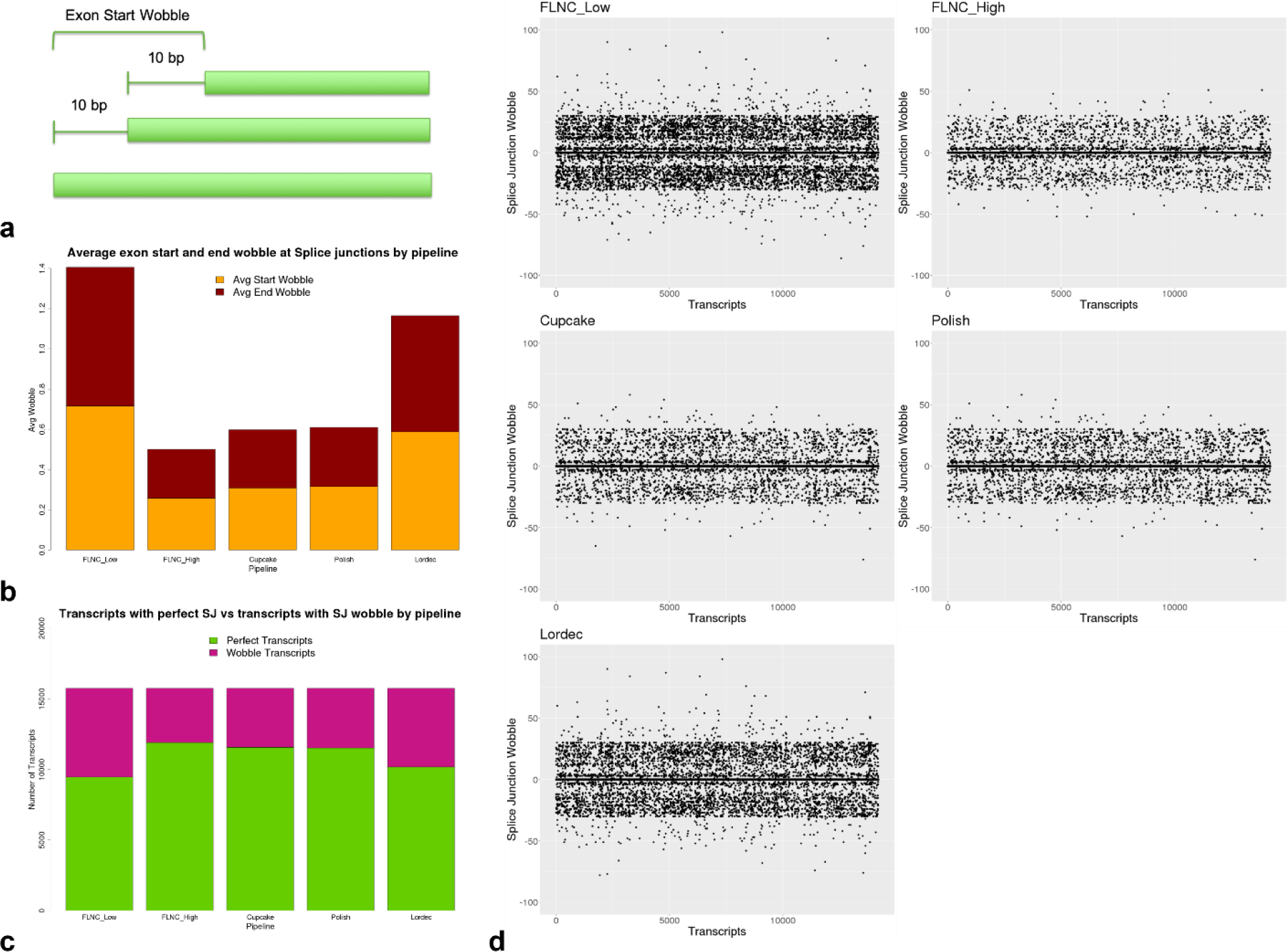
Assessing wobble across pipelines. (a) Illustration of wobble definition. (b) Average amount of wobble across splice junctions by pipeline. (c) Number of perfect transcript models and wobbly models by pipeline. (d) Scatter plot of splice junction wobble per transcript by pipeline. Wobble is shown as base pair distance from true splice junction exon start and end. Positive wobble represents exon start wobble and negative wobble represents exon end wobble. Each x-axis unit represents a single transcript model. A 30bp wobble threshold was used for the TAMA Merge run thus the apparent drop off in wobble outside this range.

**Figure 4.**
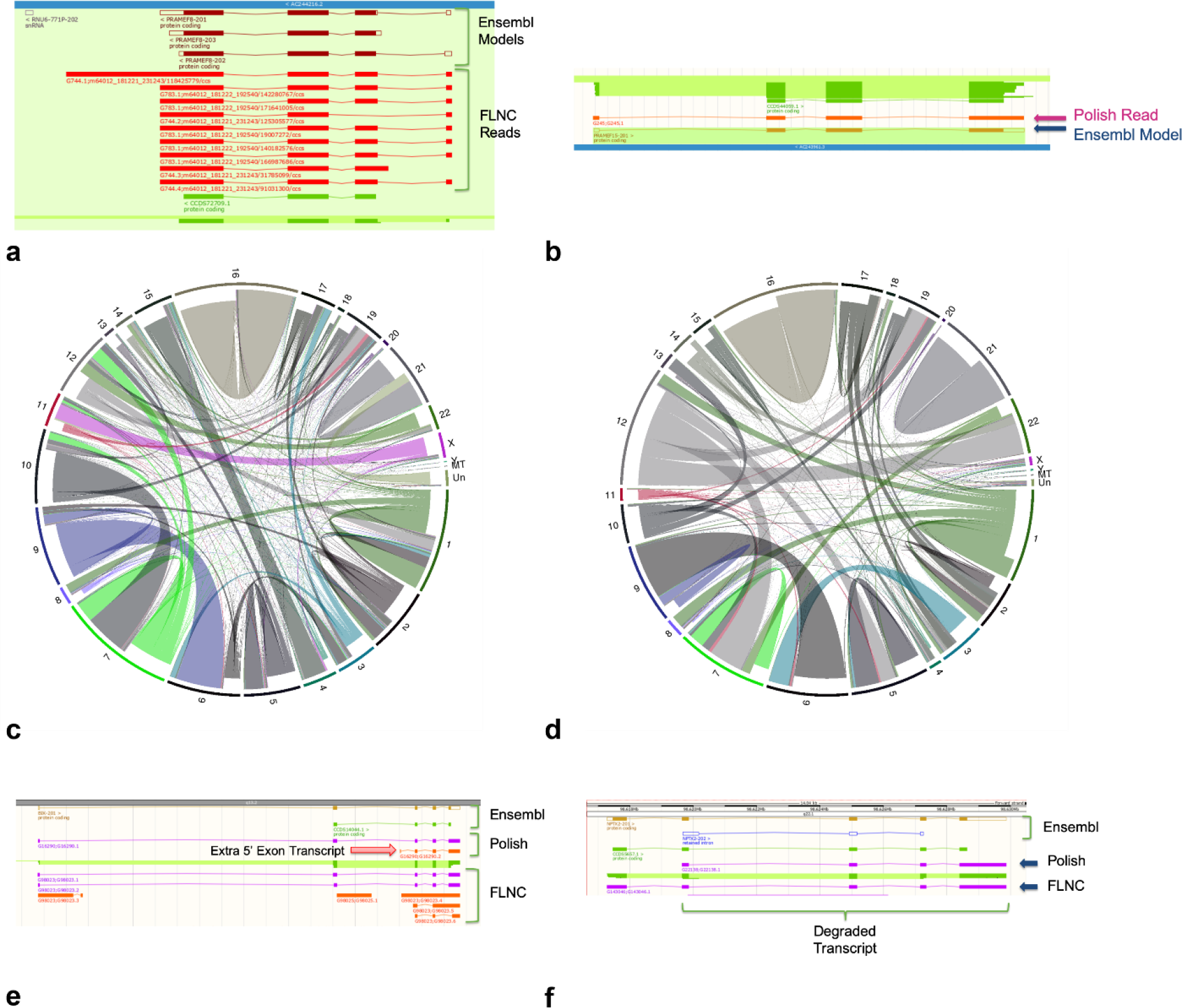
Gene and transcript read swapping from error correction. (a) PRAMEF8 gene with a coverage of 9 FLNC reads. (b) PRAMEF15 gene with a false positive coverage of one Polish read. (c) Circos plot showing reads swapping genes after correction with Cluster/Polish. Indented shows true read location and non-indented shows read allocation after error correction. Each line represents a single read moving from one gene to another with 34,637 reads from 4,799 genes moving to 2,793 genes after Cluster/Polish.(d) Circos plot for reads swapping genes after correction with Lordec. Each line represents a single read moving from one gene to another with 19,064 reads from 2,292 genes moving to 2,319 genes after Lordec error correction. (e) Exampled of false novel transcript model caused by Polish error correction. (f) Example of degraded RNA model caused by Cluster/Polish pipeline.

We compared the wobble at splice junctions for 5 different pipelines: FLNC Low, FLNC High, Polish, Cupcake, and Lordec (Fig. 3). The FLNC Low pipeline produce the highest average wobble per splice junction with 0.7155 start wobble and 0.6903 end wobble. The FLNC High pipeline had the lowest average wobble values at 0.2571 start wobble and 0.2434 end wobble. The FLNC High pipeline out performed both the Polish and Cupcake pipelines which had averages of 0.3081 start wobble and 0.2898 end wobble and 0.3149 start wobble and 0.2949 end wobble respectively. The Lordec pipeline had relatively high average wobble scores of 0.5876 start and 0.5772 end.

The FLNC Low pipeline had the lowest number of perfect transcripts (transcripts with no wobble at the splice junctions) at 9,451, while the FLNC High pipeline had the highest number of perfect transcripts at 11,891 (Fig. 3). The Polish pipeline had the second highest number of perfect transcripts at 11,562.

Thus, despite the lower overall error rates in the Polish mapped reads, the FLNC High pipeline produced a more accurate transcriptome annotation.

### Gene and transcript swapping between pipelines

One of the major concerns when using inter-read error correction methods such as Cluster/Polish and Lordec, is the possibility of creating chimeric sequences which no longer represent real transcripts. These chimeric transcripts can either be the combination of transcripts from different genes within a gene family or a combination of alternative transcripts within the same gene.

To investigate the extent to which these chimerization effects occur, we used the FLNC Low pipeline read-to-transcripts mappings as a ground truth and then looked for reads which mapped to different genes and transcripts in other pipelines. The reads that map to different loci in the Polish and Lordec pipelines represent error corrected reads that became chimeric from the inter-read correction methods. Since the Cupcake pipeline uses the same Cluster/Polish step as the Polish pipeline, there should be no differences in the read mappings. Similarly the FLNC Low and High pipelines used the same read mappings.

When comparing the FLNC Low pipeline read mappings to the Polish read mappings, there were 34,637 reads which switched from one gene locus to another after Cluster/Polish correction. A total of 6,774 genes had reads which swapped loci between the two pipelines. Of these genes, there were 3,230 genes which were only found with the FLNC Low pipeline while 104 genes were only found with the Polish pipeline. This suggests that Cluster/Polish is combining reads from different genes leading to the assignation of reads to incorrect loci.

To gain a more detailed understanding of what is happening during error correction chimerization, we examined the PReferentially expressed Antigen of MElanoma (PRAME) gene family. The PRAME gene family is highly associated with cancer development^27,28,29,30^ and is used as a biomarker for identifying various forms of cancer. Within the PRAME gene family there are 24 annotated paralogues^2^. PRAMEF8 is a gene within this family which was detected with the FLNC Low pipeline but not in the Polish pipeline. The FLNC Low pipeline shows 9 reads mapping to PRAMEF8. Of these 9 reads, 3 did not pass the Cluster/Polish step and thus were omitted from the Polish pipeline. The other 6 reads were mapped to other loci in the Polish pipeline with 5 mapping to PRAMEF15 and 1 mapping to PRAMEF27. We aligned the PRAMEF8 and PRAMEF15 transcript sequences with Muscle^31^ and found that they had 76.69% identity. While the two genes are sequentially similar, the read with the lowest identity score during genome mapping in the FLNC Low pipeline had an identity of 89.12% and 6 reads had mapping identities over 98%. Thus there is strong evidence that the reads mapped correctly in the FLNC Low pipeline and were chimerized to the point of mis-mapping in the Polish pipeline. This particular type of error could have major consequences for studies aimed at identifying gene biomarker expression.

When comparing the FLNC Low pipeline read mappings to the Lordec corrected read mappings, there were 19,064 reads which switched from one gene locus to another. A total of 3,476 genes had reads which swapped loci between the two pipelines. Of these genes there were 775 genes which were only found with the FLNC Low pipeline while 675 genes were only found with the Lordec pipeline. The number of genes found only in the Lordec pipeline is much higher than the number of genes found only in the Polish pipeline which may indicate that Lordec correction is more inclined to produce false positives.

We also examined how erroneous inter-read error correction can lead to differences in the alternative transcripts predicted. In this case, when reads from different alternative transcripts are grouped for error correction, the resulting sequence will, at best, represent only the more highly expressed transcript and, at worst, represent an artificially chimeric sequence.

Again we compared the FLNC Low pipeline to the Polish and Lordec pipelines. When comparing the FLNC Low pipeline to the Polish pipeline, we found 477,351 reads which swapped transcript models (within the same gene). This involved 112,891 transcripts with 44,852 transcripts found only in the FLNC Low annotation and 1,372 transcript found only in the Polish annotation. This represents a large difference between the two annotations with the FLNC Low pipeline predicting far more transcript models given the same reads as compared to the Polish pipeline.

When comparing the FLNC Low pipeline to the Lordec pipeline, we found 187,829 reads which swapped transcript models. This involved 142,704 transcripts with 7,117 transcripts found only in the FLNC Low annotation and 11,732 transcript found only in the Lordec annotation. Again it appears that the Lordec pipeline is more prone to producing false positives.

### Novel genes breakdown

In order to understand the source of the novel gene models found in the FLNC Low pipeline, we identified several key features of these models: coding potential, number of exons, overlap with other genes, overlap with regulatory features, and the presence of immediately downstream genomic poly-A stretches. Coding potential was assessed via three different methods: Ensembl gene match, CPAT^32^, and the TAMA open reading frame and nonsense mediated decay (TAMA ORF/NMD) pipeline. Ensembl gene matches were performed by running TAMA merge to identify gene models with overlap between the Iso-Seq annotations and the Ensembl annotation. CPAT is a tool which analyzes the sequential patterns of transcript open reading frames to predict coding potential. TAMA ORF/NMD identifies possible ORFs and then matches them to peptide sequences from the Uniprot^33^ database. If a gene did not have any predicted coding potential from any of these methods, it was labeled non-coding.

There were 26,619 gene models with coding potential and multiple exons. These are likely to represent sequences from real genes although they may not necessarily be full length models. For instance, 3,969 of these genes had downstream genomic poly-A indicating truncation of the 3’ end by the oligo-dT primer binding to the genomic poly-A region as opposed to the true poly-A tail (Fig. 5). There were 81,591 single exon non-coding genes and of those 63,919 had downstream genomic poly-A. These models may be the result of genomic fragment contamination or if the gene models overlap other genes, they may be the result of internal priming of unspliced RNA molecules.

**Figure 5.**
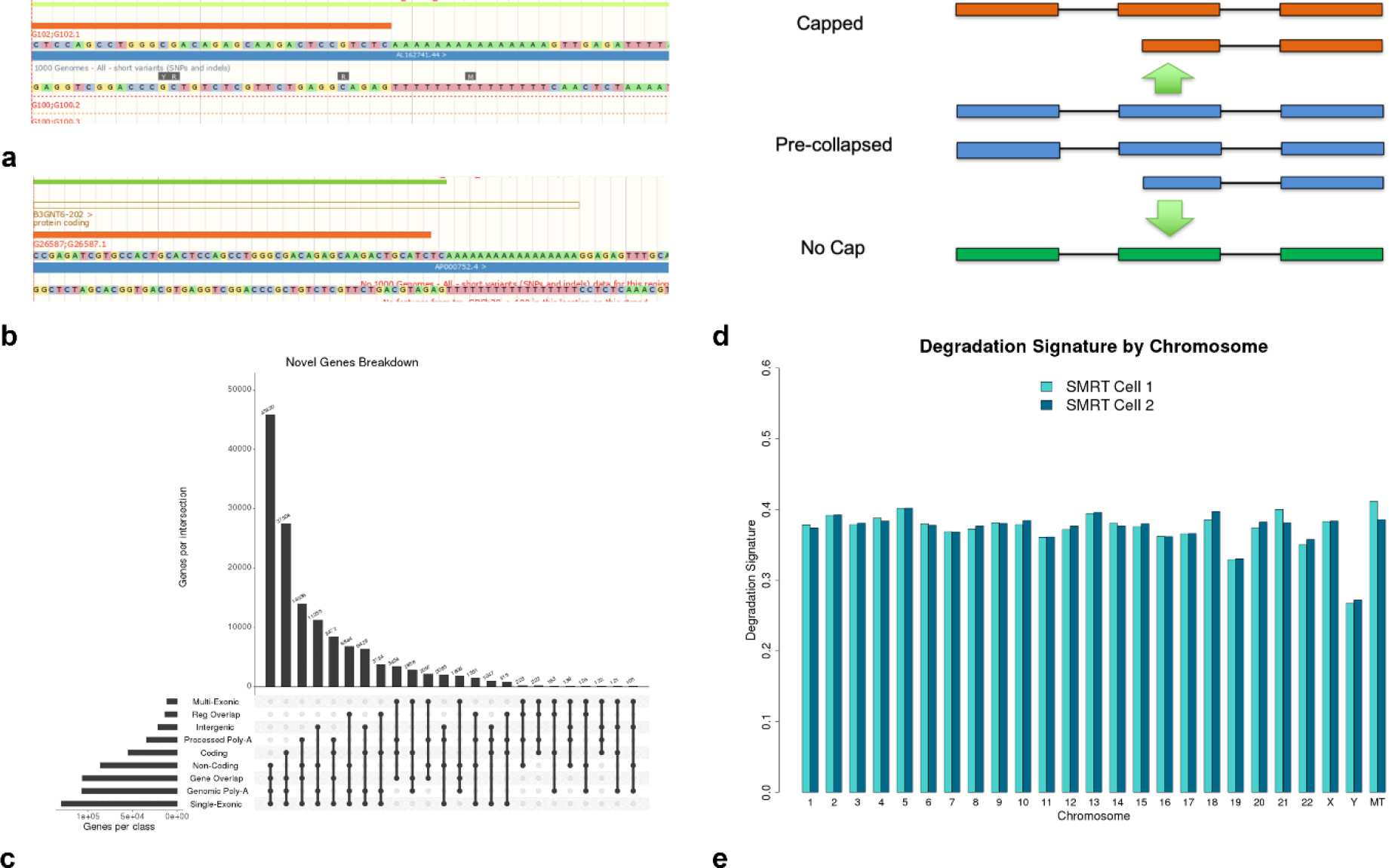
Novel genes breakdown and degradation signature analysis. (a). Example of genomic poly-A repeat within a transcribed region. Genomic poly-A can act as a site for oligo-dT primer binding thus allowing genomic fragments or internal priming to be amplified (b). Example of a real genomic poly-A repeat immediately downstream of the transcription end site (c) Novel gene breakdown by features. (d) Collapsing algorithms for 5’ cap selected RNA and non-cap selected RNA. (e). Degradation signature by chromosome per SMRT Cell run.

When looking only at the novel genes (not annotated in Ensembl), the two most common set of features are “single exonic, non-coding, gene overlap, and genomic poly-A” with 45,820 novel gene models and “single exonic, coding, gene overlap, and genomic poly-A” with 30,756 novel gene models. These are the features that gene models from internally primed un-spliced RNA would fall under. Thus it is likely that most of these models represent non-functional RNA products which make up part of the transcriptional noise of the sample.

The third most common set of features was “single exonic, non-coding, gene overlap, and processed poly-A” with 14,036 gene models. This set of features indicates the presence of real single exon gene overlapping lncRNA. Since there are no genomic poly-A stretches downstream of these models, the source RNA must have had poly-A tails added during RNA processing which would suggest the RNA truly exist in that form and may have functional roles.

### RNA Degradation

To gauge the quality of the RNA used in the Iso-Seq sequencing, we developed a metric called the Degradation Signature (DegSig) which can be calculated using the mapped reads. DegSig is defined as the percent difference in the number of transcript models for multi-read multi-exon transcripts between collapsing using a capped algorithm vs a no cap algorithm (Fig. 5). A higher percent difference indicates that the sample RNA had a greater proportion of degraded RNA.

To test the DegSig metric we used Iso-Seq data from chicken brain samples. One sample was prepared with a non-cap selecting method while the other was prepared with Teloprime^34^ 5’ cap selection. The DegSig of the non-cap selected data was 56.3% while the DegSig for the cap selected data was 23.6%. This suggests a real difference in the proportion of degraded RNA sequences captured as cDNA by the two different methods.

Since the 5’ exon cascade can exist in real gene models, we tried calculating the DegSig of the Ensembl human reference annotation to see what a reasonable baseline would be. The DegSig for the Ensembl human annotation was 1.5%. Thus a DegSig of 0% is virtually impossible, but a completely 5’ intact cDNA library could theoretically have a DegSig close to 0%. However, since models derived from 5’ degraded transcripts are difficult to distinguish from real gene models with 5’ exon cascade patterns, it may be that these types of transcript models are under-represented in all annotations which would make the estimation from the Ensembl human annotation inaccurately low.

We ran DegSig on the UHHR Iso-Seq dataset with independent calculations per SMRT cell and chromosome to see if there were any significant differences in DegSig between chromosomes.

Almost all chromosomes had a DegSig between 32% and 41% (Fig. 5). However, the Y chromosome had a DegSig of 26.7% and 27.2% for SMRT Cell 1 and 2 respectively. One explanation for the much lower DegSig on the Y chromosome may be due to the lack of read depth for the Y chromosome with only 629 and 588 reads from SMRT cells 1 and 2 respectively. Lower read depths can decrease the DegSig values due to the lack of coverage for each transcript model. For a single gene with only one transcript, there is a lower probability that 2 reads will capture a degraded transcript as compared to 100 reads.

The range of DegSig for the human data is higher than that for the chicken 5’ cap selected RNA data, thus suggesting that there may be a significant number of truncated models in the UHRR transcript annotation results. This could also be a source of the unusually high number of novel alternative transcripts predicted in the FLNC Low pipeline.

## Discussion

The UHRR PacBio Sequel II Iso-Seq dataset is the result of the most accurate long read RNA sequencing technology applied to an RNA library used as a reference for gene profiling experiments. Thus this dataset represents the technological limits and challenges that are pertinent to all RNA sequencing studies. The resulting transcriptome annotation, however, portrays a very different composition of gene models compared to public transcriptome annotations. This raises questions regarding what exactly is present in our sequencing data and what is the best way to further dissect this information to produce biologically meaningful results.

There has been a heavy emphasis on the use of multi-omics or orthogonal data to identify what is real and functional within the transcriptome. While this is certainly a powerful means of investigating novel genes, the pipelines developed for this purpose often overlook the need to properly process individual sources of data before integrating across data types. Using the TAMA tool kit, we have demonstrated some key issues with current long read RNA data pipelines that could have major effects on current transcriptomic studies.

If the goal of RNA sequencing is to accurately identify the sequences and features of real transcripts, the semantics surrounding this characterization should reflect biologically relevant information. We propose that the major features of interest for eukaryotic species should be genomic loci, transcription start and end, and splice junctions. These 4 major attributes, allows for more precise integration with other sources of information and form the backbone of defining gene models. While these features are certainly not new, the ideas pertaining to how we identify these features are still developing.

As can be seen in the difference between read errors and splice junction accuracy, one metric, although related, does not have a direct correlation with the other. Thus algorithms for achieving positional feature predictions should be designed with this foundation. While error correction is a worthy objective, it cannot be applied at the cost of biological inaccuracies as is the case for the gene and transcript swapping events occurring as a result of long read and short read error correction.

The underlying issues in all methodologies is the balance between retaining useful information and discarding misleading information. The TAMA tool kit is based around the philosophy that data should only be discarded if there is evidence that it is erroneous. This differs from other methodologies which try to preserve information which is seemingly real due to orthogonal information but may in fact still be erroneous. This philosophy limits both sensitivity and specificity for gene discovery.

From our analyses of the UHRR PacBio Sequel II Iso-Seq data, we have identified that either there are issues with the RNA preparation methods of the Universal Human Reference RNA or there are still thousands of novel genes that have not been annotated in the human genome.

## Methods

### Universal Human Reference RNA

An RNA library was first created by pooling the Universal Human Reference RNA (Agilent) with SIRV Isoform Mix E0 (Lexogen). cDNA was prepared from the RNA using the Clontech SMARTer kit. The sequencing library was the prepare using the Iso-Seq Template Preparation for Sequel Systems (PN 101-070-200) and Sequencing Sequel System II with “Early Access” binding kit (101-490-800) and chemistry (101-490-900). The sequencing library was sequenced on two Sequel II SMRT cells.

### Chicken Brain RNA

The non-cap selected chicken brain Iso-Seq data is from the European Nucleotide Archive submission PRJEB13246 which was previously analyzed and published^3^.

The cap selected chicken brain Iso-Seq data was from an adult Advanced Intercross Line chicken whole brain sample. The RNA was extracted from the tissue sample using the Qiagen RNeasy Mini Kit. The RNA was converted to cDNA using the Lexogen Teloprime kit. The resulting cDNA library was sent to Edinburgh Genomics for sequencing on the Sequel 1 system using 2.0 chemistry.

### Iso-Seq Processing

The UHRR Sequel II Iso-Seq data was processed from subread level using the CCS tool with the parameters “--noPolish --minPasses=1”. The resulting CCS reads were then stripped of adapter sequences using LIMA (lima --isoseq --dump-clips). The poly-A tails were then removed using the Refine tool (isoseq3 refine --require-polya).

### FLNC Low Pipeline

For the FLNC Low pipeline, the resulting FLNC reads were mapped to the human reference genome (GRCh38) using Minimap2 (--secondary=no -ax splice -uf -C5 -t 8). The resulting bam file was then split into 12 smaller bam files using tama_mapped_sam_splitter.py which splits bam files by chromosome thus preventing splitting between reads from the same gene. The resulting smaller bam files were then processed using TAMA Collapse (-d merge_dup -x no_cap -a 100 -z 100 -sj sj_priority – lde 5 -sjt 20 −log log_off). The resulting annotation bed files were then merged into a single bed file using TAMA Merge (-a 100 -z 100). The merged bed files from each SMRT cell were then merged together using TAMA Merge (-a 100 -z 100).

### FLNC High Pipeline

For the FLNC High pipeline, TAMA Collapse was run on the split bam files using more stringent parameters that filter out any mapped read with more than 1 error within 20 bp of a splice junction (-d merge_dup -x no_cap -a 100 -z 100 -sj sj_priority -lde 1 -sjt 20 −log log_off). The resulting annotation bed files were then merged into a single bed file using TAMA Merge (-a 100 -z 100). The merged bed files from each SMRT cell were then merged together using TAMA Merge (-a 100 -z 100). Then transcript models that were only supported by a single read were filtered out using tama_remove_single_read_models_levels.py (-l transcript -k remove_multi -s 2).

### Polish Pipeline

The resulting FLNC reads from the Refine step were clustered using Iso-Seq3 Cluster with default parameters. Iso-Seq3 Polish was then run to perform inter-read error correction. The resulting cluster reads were then mapped to the genome using Minimap2 (--secondary=no -ax splice -uf -C5 -t 8). The resulting bam file was processed using TAMA Collapse (-d merge_dup -x no_cap -a 100 -z 100 -sj sj_priority -lde 5 -sjt 20 −log log_off).

### Cupcake Pipeline

The same steps were performed as in the Polish pipeline up to mapping. After mapping, the resulting bam file was processed using Cupcake collapse_isoforms_by_sam.py (--dun-merge-5-shorter).

### Lordec Pipeline

The FLNC reads from the Isoseq3 refine step were error corrected using Lordec (-k 31 -s 3) with short read RNAseq data from the Universal Human Reference RNA (Agilent) (https://www.ncbi.nlm.nih.gov/sra/SRX1426160) (https://rnajournal.cshlp.org/content/22/4/597.full.pdf). The resulting error corrected reads were then processed in the same way as the FLNC Low starting from mapping to the genome.

### Wobble Analysis

To assess the wobble between each pipeline and the Ensembl annotation, we used TAMA merge with parameter settings (-a 300 -z 300 -m 30 -d merge_dup) which considers any transcripts which have up to 300 bp difference in their transcription start and end and up to 30 bp difference in their splice junctions starts and ends to have “nearly identical structures”. This is the definition for matching at transcript level.

### Coding Potential Analysis

For the Ensembl match evidence of coding potential, we labelled the Iso-Seq annotation genes as coding if they had any overlap on the same strand as an Ensembl annotated protein coding gene.

CPAT was used with default parameters and the built-in Human Hex models. A cutoff score of 0.364 was used to segregate between coding and non-coding transcripts.

In the ORF/NMD pipeline, ORF’s were predicted from each transcript sequence. The ORF’s were then translated into amino acid sequences. Blastp (-evalue 1e-10 – ungapped -comp_based_stats F) was used to match the amino acid sequences with the UniRef90 database. The top hits were used to select the best ORF prediction for each transcript model. Transcripts with no hits were considered to have no coding evidence from this analysis.

## Code Availability

TAMA is available from https://github.com/GenomeRIK/tama.

## Data Availability

The PacBio Universal Human Reference RNA Sequel II Iso-Seq dataset is available from https://github.com/PacificBiosciences/DevNet/wiki/Sequel-II-System-Data-Release:-Universal-Human-Reference-(UHR)-Iso-Seq. The short read Illumina RNAseq data used for Lordec error correction are available in the National Center for Biotechnology Information Sequence Read Archive under accession number SRP066009 (https://www.ncbi.nlm.nih.gov/sra/SRX1426160). The non-cap selected chicken brain Iso-Seq data is available from the European Nucleotide Archive under accession number PRJEB13246. The Teloprime cap selected chicken brain Iso-Seq data is available from the European Nucleotide Archive under accession number PRJEB25416.

## Author’s contributions

RIK developed TAMA and implemented the different Iso-Seq pipelines. RIK, DWB, and YC conceived the idea of this study. DWB provided guidance on the focus of the study. YC ran the ORF/NMD pipeline and identified issues with gene swapping. JS and ALA reviewed and edited the manuscript.

## Acknowledgements

We would like to thank Dr. Elizabeth Tseng and Pacific Biosciences for releasing the Universal Human Reference RNA Sequel II Iso-Seq dataset and providing guidance on the analyses.

## Conflict of Interest

The authors declare that they have no competing interests.

## Funding

We acknowledge funding support from the UK’s Biotechnology and Biological Sciences Research Council (Institute Strategic Programme grant BBS/E/D/10002070; and BB/N019202/1, BB/M011461/1, BB/M01844X/1). The funding bodies did not contribute to the design of the study, sample collection, analysis, interpretation of data, or in writing the manuscript.

## References

1. Harrow, J. et al. GENCODE : The reference human genome annotation for The ENCODE Project. Genome Res. 22, 1760–1774 (2012).

2. Cunningham, F. et al. Ensembl 2015. Nucleic Acids Res. 43, 662–669 (2014).

3. Kuo, R. I. et al. Normalized long read RNA sequencing in chicken reveals transcriptome complexity similar to human. BMC Genomics 18, 1–19 (2017).

4. Magrini, V. et al. Improving eukaryotic genome annotation using single molecule mRNA sequencing. 1–14 (2018). doi:10.1186/s12864-018-4555-7

5. Wang, B. et al. Unveiling the complexity of the maize transcriptome by single-molecule long-read sequencing. Nat. Commun. 7, 11708 (2016).

6. Abdel-Ghany, S. E. et al. A survey of the sorghum transcriptome using single-molecule long reads. Nat. Commun. 7, 11706 (2016).

7. Gupta, I. et al. Single-cell isoform RNA sequencing characterizes isoforms in thousands of cerebellar cells. Nat. Biotechnol. 36, 1197–1202 (2018).

8. Mays, A. D. et al. Single molecule real time (SMRT) full length RNA-sequencing reveals novel and distinct mRNA isoforms in human bone marrow cell subpopulations. 1–18 (2019). doi:10.1101/555326

9. Tseng, E. et al. Transcriptional fates of human-specific segmental duplications in brain. Genome Res. 28, 1566–1576 (2018).

10. Gao, Y. et al. PRAPI: post-transcriptional regulation analysis pipeline for Iso-Seq. Bioinformatics 34, 1580–1582 (2017).

11. Wyman, D. et al. A technology-agnostic long-read analysis pipeline for transcriptome discovery and quantification. (2019).

12. Tardaguila, M. et al. SQANTI: extensive characterization of long read transcript sequences for quality control in full-length transcriptome identification and quantification. Doi.Org 118083 (2017). doi:10.1101/118083

13. Salmela, L. & Rivals, E. LoRDEC: Accurate and efficient long read error correction. Bioinformatics 30, 3506–3514 (2014).

14. Hu, R., Sun, G. & Sun, X. LSCplus: A fast solution for improving long read accuracy by short read alignment. BMC Bioinformatics 17, 1–9 (2016).

15. Gordon, S. P. et al. Widespread polycistronic transcripts in fungi revealed by single-molecule mRNA sequencing. PLoS One 10, 1–15 (2015).

16. Hackl, T., Hedrich, R., Schultz, J. & Förster, F. Proovread: Large-scale high-accuracy PacBio correction through iterative short read consensus. Bioinformatics 30, 3004–3011 (2014).

17. Giuffra, E. & Tuggle, C. K. Functional Annotation of Animal Genomes (FAANG): Current Achievements and Roadmap. Annu. Rev. Anim. Biosci. 7, 65–88 (2019).

18. Koepfli, K.-P., Paten, B. & O’Brien, S. J. The Genome 10K Project: A Way Forward. Annu. Rev. Anim. Biosci. 3, 57–111 (2015).

19. Zerbino, D. R. et al. Ensembl 2018. Nucleic Acids Res. 46, D754–D761 (2018).

20. S.A. Zahler and J.M. Calvo. Charging it cancels the order In a newly discovered mechanism for regulating transcription termination, a. Curr. Biol. 4, 73–75 (1994).

21. Sessegolo, C. et al. Transcriptome profiling of mouse samples using nanopore sequencing of cDNA and RNA molecules. bioRxiv 575142 (2019). doi:10.1101/575142

22. Curwen, V. et al. The Ensembl automatic gene annotation system. Genome Res. 14, 942–50 (2004).

23. Nottingham, R. M. et al. RNA-seq of human reference RNA samples using a thermostable group II intron reverse transcriptase. Rna 22, 597–613 (2016).

24. Holmes, I. & Durbin, R. Dynamic programming alignment accuracy. J. Comput. Biol. 5, 493–504 (1998).

25. Schor, I. E. et al. Promoter shape varies across populations and affects promoter evolution and expression noise. Nat. Genet. (2017). doi:10.1038/ng.3791

26. Djebali, S. et al. Landscape of transcription in human cells. Nature 489, 101–8 (2012).

27. Epping, M. T., Hart, A. A. M., Glas, A. M., Krijgsman, O. & Bernards, R. PRAME expression and clinical outcome of breast cancer. Br. J. Cancer 99, 398–403 (2008).

28. Field, M. G. et al. PRAME as an independent biomarker for metastasis in uveal melanoma. Clin. Cancer Res. 22, 1234–1242 (2016).

29. Roszik, J. et al. Overexpressed PRAME is a potential immunotherapy target in sarcoma subtypes. Clin. Sarcoma Res. 7, 1–7 (2017).

30. Zhang, W. et al. PRAME expression and promoter hypomethylation in epithelial ovarian cancer. Oncotarget 7, (2016).

31. Edgar, R. C. MUSCLE: Multiple sequence alignment with high accuracy and high throughput. Nucleic Acids Res. 32, 1792–1797 (2004).

32. Wang, L. et al. CPAT: Coding-Potential Assessment Tool using an alignment-free logistic regression model. Nucleic Acids Res. 41, e74 (2013).

33. The UniProt Consortium. UniProt: a hub for protein information. Nucleic Acids Res. 43, D204–12 (2014).

34. Cartolano, M., Huettel, B., Hartwig, B., Reinhardt, R. & Schneeberger, K. cDNA library enrichment of full length transcripts for SMRT long read sequencing. PLoS One 11, 1–10 (2016).

